# Dichotomous Intrinsic Properties of Adult Accumbens Medium Spiny Neurons Vanish in the Fragile X Mouse Model of Autism

**DOI:** 10.1101/2022.07.06.498928

**Authors:** Gabriele Giua, Olivier Lassalle, Leila Makrini-Maleville, Emmanuel Valjent, Pascale Chavis, Olivier J.J. Manzoni

## Abstract

Fragile X syndrome (FXS), the most common cause of autism and inherited intellectual disability, is caused by the mutation of a single gene, *fmr1*, which encodes the Fragile X mental retardation protein (FMRP). FXS patients suffer from cognitive, emotional, and social deficits indicative of dysfunction in the nucleus accumbens (NAc), a structure central to the control of social behavior. The major cell type of the NAc, medium spiny neurons (MSNs), are differentiated in two subtypes based on their expression of either dopamine D1 or D2 receptors, their connectivity, and associated behavioral functions. Understanding how the absence of FMRP differentially affects the cellular properties of MSNs is a necessary step to categorize FXS cellular endophenotypes. To address this question, we comprehensively compared the intrinsic passive and active properties of MSN subtypes identified in a novel *Fmr1-/y* :: *Drd1a-tdTomato* mouse model allowing in-situ identification of MSN subtypes in FXS mice. Although *fmr1* transcripts and their gene product, FMRP, were found in both MSNs subtypes, the results suggest cell-autonomous functions for *Fmr1*. The opposite membrane properties and action potential kinetics that normally discriminate D1- from D2- MSNs in WT mouse is either reversed or abolished in *Fmr1-/y* :: *Drd1a-tdTomato* mice. Multivariate analysis shed light on the compound effects of *Fmr1* ablation by revealing how the phenotypic traits that distinguish each cell type in WT are modified in FXS. Together these data show that in Fragile X mice the normal dichotomy that characterizes NAc D1- and D2-MSNs is thrown out of balance, leading to a uniform phenotype that could underlie selected aspects of the pathology.

## Background

Fragile X Syndrome (FXS) is caused by meiotic instability in the 5’ untranslated region of the X chromosome-linked fragile X mental retardation 1 gene (*Fmr1*), impeding the transcription of the Fragile X Mental Retardation protein (FMRP) [1]. FMRP is an RNA-binding protein, widely expressed in the central nervous system, which regulates the translation of thousands of mRNA targets and thus its loss in FXS largely modifies protein synthesis [2–4]. Among the pleiotropic outcomes of FMRP absence, altered neuronal and circuit excitability is thought to be a major component of FXS phenotypes [5]. The variety of neuropsychiatric symptoms in FXS, such as learning disabilities, hyperactivity, repetitive behaviors, impulsivity, and symptoms affecting social behaviors associated with autism spectrum disorder, suggests dysfunction in multiple brain areas. Cortical [6–13] and hippocampal [14] dysfunctions have been extensively studied. Although the symptomology of FXS includes social deficits [15] compatible with altered neuronal functions in the mesolimbic network and the basal ganglia [16,17], studies addressing how FXS modifies neuronal excitability in these systems are scarce. The central part of the mesolimbic pathway, the nucleus accumbens (NAc), processes emotional, motivational, reward and aversion signals and plays a role in behaviors essential to the survival of the animal [18,19]. The lack of FMRP modifies synaptic long-term potentiation [20], long-term depression [21,22] in the NAc as well as the dendritic morphology of the principal cell-type of the NAc, medium spiny neurons (MSNs) [20]. NAc MSNs are GABAergic projection neurons which express either D1 or D2 receptors (D1R or D2R) and play specific roles in NAc-mediated behaviors and disorders [23,24]. Several common symptoms of autism spectrum disorders observed in FXS indicate an altered balance between approach and avoidance responses. Specifically, direct pathway D1-MSNs mediate approach behavioral responses while D2-MSNs mediate avoidance behavioral responses via the so-called indirect pathway [17]. Here we explored the cell-type specific intrinsic properties of MSNs in the NAc Core of a novel *Fmr1+/y* or *-/y* :: *Drd1a-tdTomato* mouse model allowing in-situ identification of MSN subtypes in WT and FXS littermate mice. Overall, the results show the cell-specific endophenotypes of FMRP’s ablation in the NAc: the normal dichotomy that characterizes D1- and D2-MSNs is thrown out of balance, leading to a uniform phenotype that could underlie selected aspects of the pathology.

## Materials and Methods

### Animals

Animals were treated in compliance with the European Communities Council Directive (86/609/EEC) and the United States National Institutes of Health Guide for the care and use of laboratory animals. The French Ethical committee authorized this project (APAFIS#3279-2015121715284829 v5). Three different cohorts of mice were used in this study. (*1*) Mice implied in electrophysiological experiments were obtained breeding *Drd1a-tdTomato* x *Fmr1* KO2 mice from the Jackson Laboratory (Bar Harbor, ME, USA) and FRAXA foundation respectively. Both strains had a C57Bl/6J background. Mice were acclimated to the animal facility for one week and then housed in male *Drd1a-tdTomato* and female *Fmr1*+/- pairs for breeding. Pups were weaned and ear punched for identification and genotyping at P21. Ex-vivo electrophysiological recordings were performed on first-generation male mice between P70 and P100. *Fmr1+/y* :: *Drd1a-tdTomato* mice composed the control group (wild type (WT)) and *Fmr1-/y* :: *Drd1a-tdTomato* the experimental group (knockout (KO)). (*2*) For immunofluorescence experiments we used a cohort of 8–12-week-old male mice *Drd1a*-*EGFP* (enhanced green fluorescent protein) (n = 3, C57BL/6 background, founder *S118*), generated by GENSAT (Gene Expression Nervous System Atlas, Rockefeller University, New York, NY, USA). (*3*) Male C57BL/6J mice 8 weeks old from Jackson Laboratory were used for in situ hybridization (ISH) assay (n=2). All mice used in this study were housed in groups of 4-5 mice at constant room temperature (20 ± 1°C) and humidity (60%) and exposed to a light cycle of 12h light/dark with ad libitum access to food and water.

### Immunofluorescence assay

Mice were rapidly anaesthetized with Euthasol (360 mg/kg, i.p., TVM lab, France) and transcardially perfused with 4% (weight/vol) paraformaldehyde in 0.1 M sodium phosphate buffer (PBS) (pH 7.5) [27]. Brains were post-fixed overnight at 4°C in the same solution. Thirty-μm thick sections were cut with a vibratome (Leica, France) and stored at −20°C in a solution containing 30% (vol/vol) ethylene glycol, 30% (vol/vol) glycerol, and 0.1M sodium phosphate buffer, until they were processed for immunofluorescence. NAc Core sections were identified using a mouse brain atlas and sections comprised between 1.54 mm and 1.42 mm from bregma were included in the analysis. Free-floating sections were rinsed three times 10 min in PBS followed by an incubation of 15 min in 0.2% (vol/vol) Triton X-100 in PBS. Sections were rinsed again in PBS and blocked for 1 hour in a solution of 3% BSA in PBS, before being incubated 72hours at 4°C with the following primary antibodies: chicken anti-GFP (1:1000, Life Technologies, #A10262) and rabbit anti-FMRP (1:500, Millipore, #ab60-46). Sections were rinsed three times for 10 min in PBS and incubated for 45 min with goat Cy3-coupled anti-rabbit (1:500, Thermo Fisher Scientific Cat# 10520) and goat Alexa Fluor 488-coupled anti-chicken (1:500, Thermo Fisher Scientific Cat#A-11039). Sections were rinsed for 10 min twice in PBS before mounting in DPX (Sigma-Aldrich).

### RNAscope ISH assay

Staining for *Drd1* (dopamine receptor D1 gene), *Drd2* (dopamine receptor D2 gene) and *Fmr1* mRNAs was performed using single molecule fluorescent in situ hybridization (smFISH). Brains from 2 C57BL/6J (8 weeks old) male mice were rapidly extracted and snap-frozen on dry ice and stored at -80°C until use. Ventral striatum coronal sections (14 μm) were collected directly onto Superfrost Plus slides (Fisherbrand). RNAscope Fluorescent Multiplex labeling kit (ACDBio Cat No. 320850) was used to perform the smFISH assay according to manufacturer’s recommendations. Probes used for staining are mm-Drd1-C1 (ACDBio Cat No. 461901), mm-Drd2-C3 (ACDBio Cat No. 406501-C3) and mm-Fmr1-C2 (ACDBio Cat No. 496391-C2). After incubation with fluorescent-labeled probes, slides were counterstained with DAPI and mounted with ProLong Diamond Antifade mounting medium (Thermo Fisher scientific P36961).

### Images acquisition

Confocal microscopy and image analysis were carried out at the Montpellier RIO Imaging Facility. Images were acquired using sequential laser scanning confocal microscopy (Leica SP8). Photomicrographs were obtained with the following band-pass and long-pass filter setting: Alexa Fluor 488/Cy2 (band pass filter: 505–530), Cy3 (band pass filter: 560–615) and Cy5 (long-pass filter 650). All parameters were held constant for all sections from the same experiment.

### Slice preparation for ex-vivo electrophysiological recordings

Adult male mice (P70-P100) were deeply anesthetized with isoflurane and decapitated according to institutional regulations, as previously described [25]. The brain was sliced (300 μm) on the coronal plane with a vibratome (Integraslice, Campden Instruments) in a sucrose-based solution at 4°C (NaCl 87mM, sucrose 75mM, glucose 25mM, KCl 2.5mM, MgCl_2_ 4mM, CaCl_2_ 0.5mM, NaHCO_3_ 23mM and NaH2PO_4_ 1.25mM). Immediately after cutting, slices containing the NAc Core were stored for 1 h at 32°C in a low calcium artificial cerebrospinal fluid (ACSF; NaCl 130mM, glucose 11mM, KCl 2.5mM, MgCl_2_ 2.4mM, CaCl_2_ 1.2mM, NaHCO_3_ 23mM and NaH_2_PO_4_ 1.2mM), equilibrated with 95% O_2_/5% CO_2_. After 1 hour of recovery, slices were kept at room temperature until the time of recording.

### Electrophysiology

Whole-cell patch-clamp recordings were collected from MSNs of NAc Core. MSNs were visualized using an upright microscope with infrared illumination and then distinguished in D1 or D2 expressing MSNs based on the visualization of *Drd1*-tdTomato using an upright microscope with infrared and fluorescent illumination. During the recording, coronal slices containing the NAc were placed in the recording chamber and superfused at 2 mL/min with normal calcium ACSF (NaCl 130mM, glucose 11mM, KCl 2.5mM, MgCl_2_ 1.2mM, CaCl_2_ 2.4mM, NaHCO_3_ 23mM and NaH_2_PO_4_ 1.2mM), equilibrated with 95% O_2_/5% CO_2_. The intracellular solution was based on K^+^ gluconate (K^+^ gluconate 145mM, NaCl 3mM, MgCl_2_ 1mM, EGTA 1mM, CaCl_2_ 0.3mM, Na_2_ATP 2mM, NaGTP 0.5mM, cAMP 0.2mM, buffered with HEPES 10mM). Its pH was adjusted to 7.2 and osmolarity to 290-300mOsm. Electrode resistance was 3-5 MOhm. All experiments were done at 25±1°C. The superfusion medium contained gabazine 10μM (SR 95531 hydrobromide; Tocris) to block GABA Type A (GABA-A) receptors.

Data was recorded in current clamp with an Axopatch-200B amplifier, low pass filtered at 2 kHz, digitized (10 kHz, DigiData 1440A, Axon Instruments), collected and analyzed using Clampex 10.7 (Molecular Device).

MSNs intrinsic properties were obtained from current-voltage (I-V) curves. I-V curves were made by a series of hyperpolarizing to depolarizing current steps (500ms) immediately after breaking into the cell.

### Statistical analysis

Data were tested for Normality (D’Agostino-Pearson and Shapiro-Wilk tests) and outliers (ROUT test) before statistical analysis. Nonparametric Mann-Whitney *U* and Spearman correlation tests were performed with Prism (GraphPad Software). Correlations were compared with “cocor.indep.groups” of the package cocor [26] in R 4.2.0 (RCore Team 2020). Comparison results were shown only for those correlations found to be significant in at least one of the two compared groups. The index was built with input resistance, APs number at +150pA current step and AP duration, which reflect passive membrane properties, neuronal firing properties and APs kinetics, respectively. The values were standardized for each variable by scaling (mean of 0 and SD of 1) and pooled for each group. N values represent individual animals. Statistical significance was set at p < 0.05.

## Results

The intrinsic properties of identified D1- and D2-MSNs in the NAc Core of *Fmr1* KO mice and WT littermates were recorded and compared in a total of 156 MSNs from 49 young adult *Fmr1*+/y or -/y :: *Drd1a-tdTomato* male mice (P70-P100). The dataset was divided into four groups: tdTomato-labeled *Drd1a*-positive MSNs from WT mice (WT D1), tdTomato-unlabeled MSNs from WT mice (WT D2), tdTomato-labeled *Drd1a*-positive MSNs from *Fmr1* KO mice (KO D1), tdTomato-unlabeled MSNs from *Fmr1* KO mice (KO D2). We and others have consistently shown that unlabeled D1-negative MSNs are all D2-positive MSNs, thus we refer to tdTomato-unlabeled MSNs as D2 MSNs [25,28–32]. The analysis was designed to highlight: (*i)* differences between MSN subtypes within each genotype; (*ii*) differences between the two genotypes for the same MSN subtype.

### *Fmr1* mRNA is expressed in both D1- and D2-MSNs in the NAc Core of WT mice

First, the expression of *Fmr1* transcripts was quantified in a dataset of RNAseq generated on tagged ribosome-bound mRNAs and the input fractions of NAc extract (including both Core and shell) of D2-Ribotag and Wfs1-Ribotag mice [33]. *Fmr1* transcripts were less abundant (adjusted P value p < 0.05) in the NAc pellet fraction of D2-RiboTag mice compared with the input fraction (containing the mRNAs from all cellular types) suggesting that *Fmr1* gene products were less expressed in D2-MSNs (**Fig. 1A, left**). These results were in line with the slight enrichment of *Fmr1* transcripts in the NAc pellet fraction of Wfs1-MSNs, a NAc Core MSNs population enriched in *Drd1* transcripts [33] (**Fig. 1A, right**). The presence of *Frm1* mRNA in NAc Core MSNs was confirmed by *in situ* hybridization. Indeed, *Fmr1* transcripts were found in striatal cells expressing the gene encoding for the dopamine D1 and D2 receptors (*Drd1* and *Drd2*), identifying D1- and D2-MSNs, respectively (**Fig. 1B**). The distribution of FMRP in both MSNs population was finally validated at the protein level. Double immunofluorescence analysis of the NAc Core of Drd1-EGFP mice revealed that FMRP was detected in both GFP-positive (D1-MSNs) and GFP-negative (D2-MSNs) [28] **(Fig. 1C)**. Together, these results indicate that both NAc Core D1- and D2-MSNs express FMRP.

**Figure 1.**
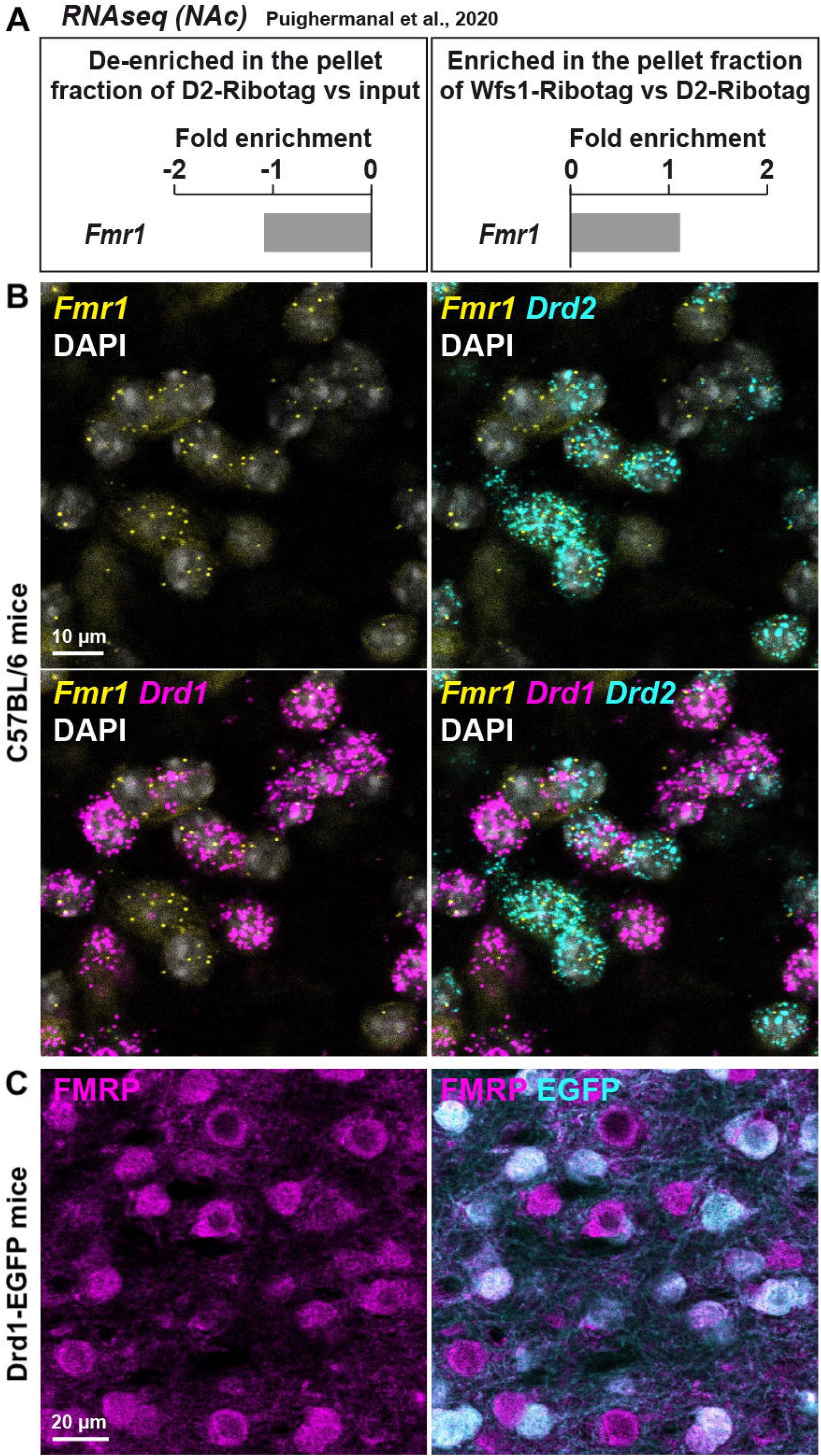
*Fmr1* mRNA is detected in both D1- and D2-MSNs in the NAc Core of WT mice. **a**) Fold-change of *Fmr1* transcripts found by RNAseq in NAc extract of D2-Ribotag and Wfs1-Ribotag mice (Puighermanal et al., 2020). *Fmr1* transcripts were slightly “de-enriched” in the NAc pellet fraction of *D2-RiboTag* mice compared with the input fraction (containing the mRNAs from all cellular types) (lower panel) but slightly enriched in the NAc pellet fraction of *Wfs1-RiboTag* mice (right panel). **b**) Single-molecular fluorescent in situ hybridization for *Drd1* (magenta), *Drd2* (cyan) and *Fmr1* (yellow) mRNAs in the NAc Core. Slides were counterstained with DAPI (white). Scale bar: 10 µm. *Fmr1* mRNA expression is detected in both *Drd1*-positive and *Drd2*-positive neurons. **c**) Double immunofluorescence for FMRP (magenta) and EGFP (cyan) in the NAc Core of *Drd1*-EGFP mice. Scale bars: 20 μm.

### Discrepancies in the passive properties of accumbal D1- and D2-MSNs in WTs vanishes in FXS mice

The passive properties of adult current-clamped and visually-identified neighboring D1 and D2 MSNs were compared in the NAc Core of WT and *Fmr1* KO littermates **(Fig. 2, Suppl. Fig. 1)**. In WT mice, the resting potential of D1- and D2-MSNs is similar. In contrast, D2-MSNs were more depolarized than D1-MSNs in *Fmr1* KO mice **(Fig. 2B)**. No differences in the firing latency were observed in WTs while in *Fmr1* KO mice D2-MSNs fired more readily than D1-MSNs **(Fig. 2C)**. In WT mice, D2-MSNs displayed a higher rheobase compared to D1-MSNs whereas in *Fmr1* KO this difference disappeared. Here, D2-MSNs of *Fmr1* KO mice showed a lower rheobase when compared to D2-MSNs of WTs, according to their depolarized state **(Fig. 2E)**. Moreover, D2-MSNs exhibited a more negative threshold than D1-MSNs in WTs and this dichotomy vanished in *Fmr1* KO mice. Differentiating cell subtypes showed that only D1-MSNs varied between genotypes, with a more negative threshold in *Fmr1* KO mice **(Fig. 2F)**. As shown in **Fig. 2G**, the membrane voltage responses to hyperpolarizing current steps strongly differed between D1- and D2-MSNs in WTs **(Fig. 2H)** but were similar in *Fmr1* KO mice **(Fig. 2I)**. This difference likely reflects an alteration in input resistance in the absence of FMRP **(Fig. 2J, K; Suppl. Fig. 1D, E)**. Specifically, in *Fmr1* KO mice the input resistance of D1-MSNs was decreased **(Suppl. Fig. 1D)** whereas that of D2-MSNs was increased **(Suppl. Fig. 1E)** when compared with WT. Finally, the membrane response to a somatic injection of -400pA also revealed a higher sag current in the D2-MSNs of *Fmr1* KO mice when compared with those of WTs **(Suppl. Fig. 1G)**.

**Figure 2.**
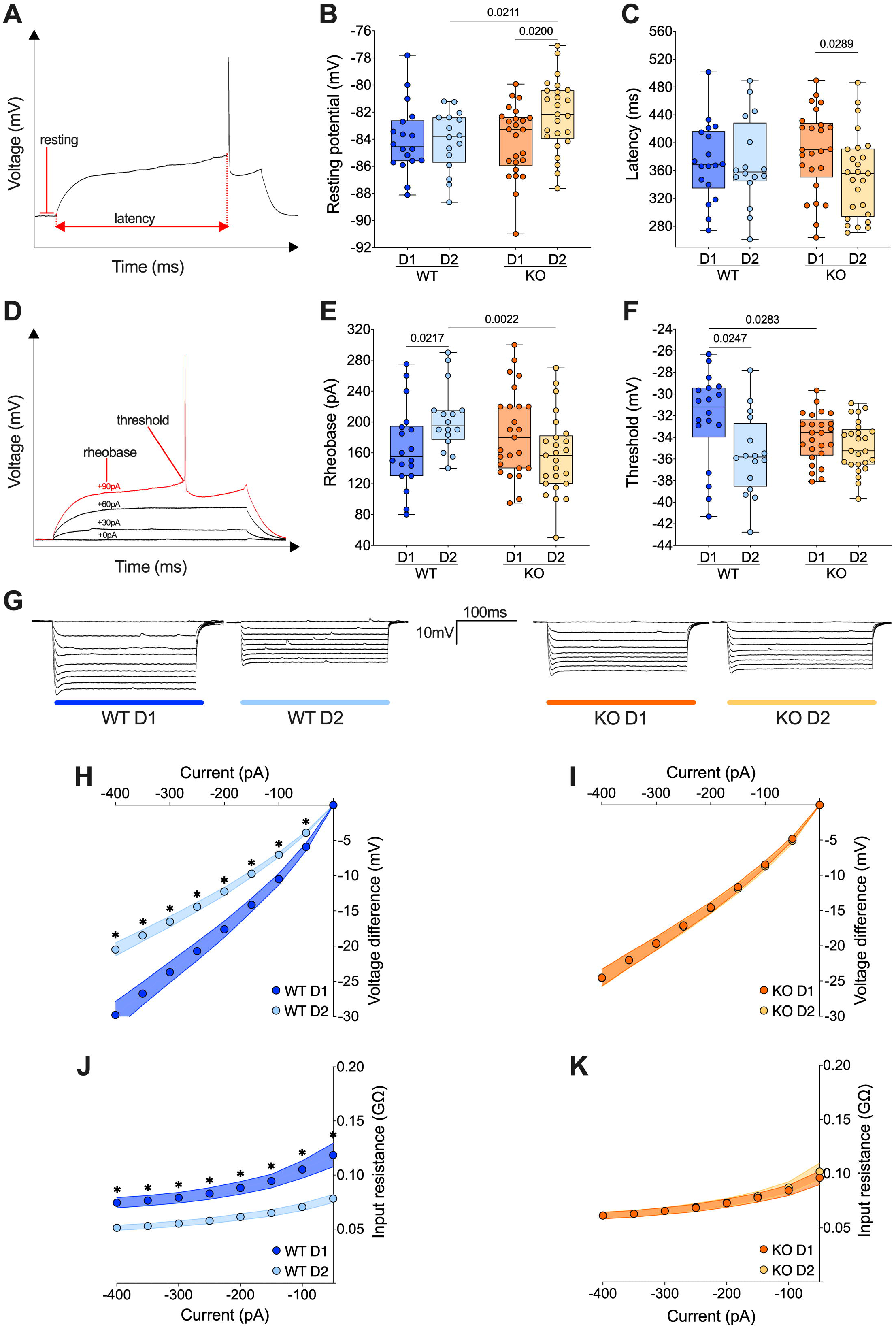
The normal discrepancy in the passive properties of accumbal D1- and D2- MSNs in wild type vanishes in FXS mice. **(A)** Example of individual AP evoked by depolarizing current injection, indicating resting potential and latency metrics. **(B)** The resting membrane potential of D2-MSNs is significantly depolarized in *Fmr1* KO mice compared to that of D2-MSNs of WT mice and D1-MSNs recorded in *Fmr1* KOs. **(C)** In *Fmr1* KO mice D2-MSNs showed shorter firing latency than D1-MSNs. **(D)** Example of individual AP evoked by increasing current steps, indicating rheobase and action potential threshold. **(E)** In WT mice the rheobase is higher in D2-MSNs than in D1-MSNs but in *Fmr1* KO mice the rheobases are similar. **(F)** In WT mice, D1-MSNs have a higher spiking threshold than D2-MSNs. In contrast, this difference is absent in *Fmr1* KO. **(B, C, E, F)** Single dot represents an individual mouse. Data are shown as min. to max. box plot with median and 25-75 percentile. Mann-Whitney *U* tests. P-values <0.05 are displayed in graphs. **(G)** Examples of membrane voltage response to hyperpolarizing current injections in our experimental groups. **(H)** Current injection steps of 50 pA from -400 pA to -50 pA revealed differences in the I–V relationship in D1- and D2-MSNs in the accumbens of WT mice (i.e., in response to hyperpolarizing current injections, in WT mice D1-MSNs showed a greater response compared to D2-MSNs), **(I)** but not *Fmr1* KO mice. **(J)** The membrane input resistance was higher in D1-MSNs than in D2-MSNs in WT, **(K)** but not *Fmr1* KO mice. **(H-K)** Single dot represents group mean value at that current step. Data are shown as mean ± SEM in XY plot. Multiple Mann-Whitney *U* test, * p-values <0.05. **(B, C, E, F, H-K)** WT D1-MSNs N=18 in dark blue, WT D2-MSNs N=16 in light blue, KO D1-MSNs N=25 in dark orange, KO D2-MSNs N=25 in light orange.

Taken together, these results show that the profound functional differences in passive properties that normally differentiate D1-from D2-MSNs are abolished in mice lacking FMRP.

### The active properties of accumbal D1- and D2-MSNs diverge in opposite ways in WT and FXS mice

The current changes in passive properties strongly suggest cell-subtype specific modifications of MSNs’ intrinsic excitability. The membrane reaction profiles of D1- and D2-MSNs in response to a series of increasing somatic current steps differ strongly between the two genotypes **(Fig. 3A-C)**. Within physiological range, D1- were more excitable than D2- MSNs in WT mice, whereas in *Fmr1* KO mice it was the opposite **(Fig. 3A-D)**. Conversely, at higher current steps, firing accommodation occurred at lower current steps in D1- than in D2- MSNs in WTs, contrary to KO **(Fig. 3E)**. The comparison of firing profile by cell types showed that the absence of FMRP impacted the excitability of both neuronal subtypes in opposite manners **(Suppl. Fig. 2A-C)**, in parallel with opposite changes in input resistance **(Suppl. Fig. 1D, E)**.

**Figure 3.**
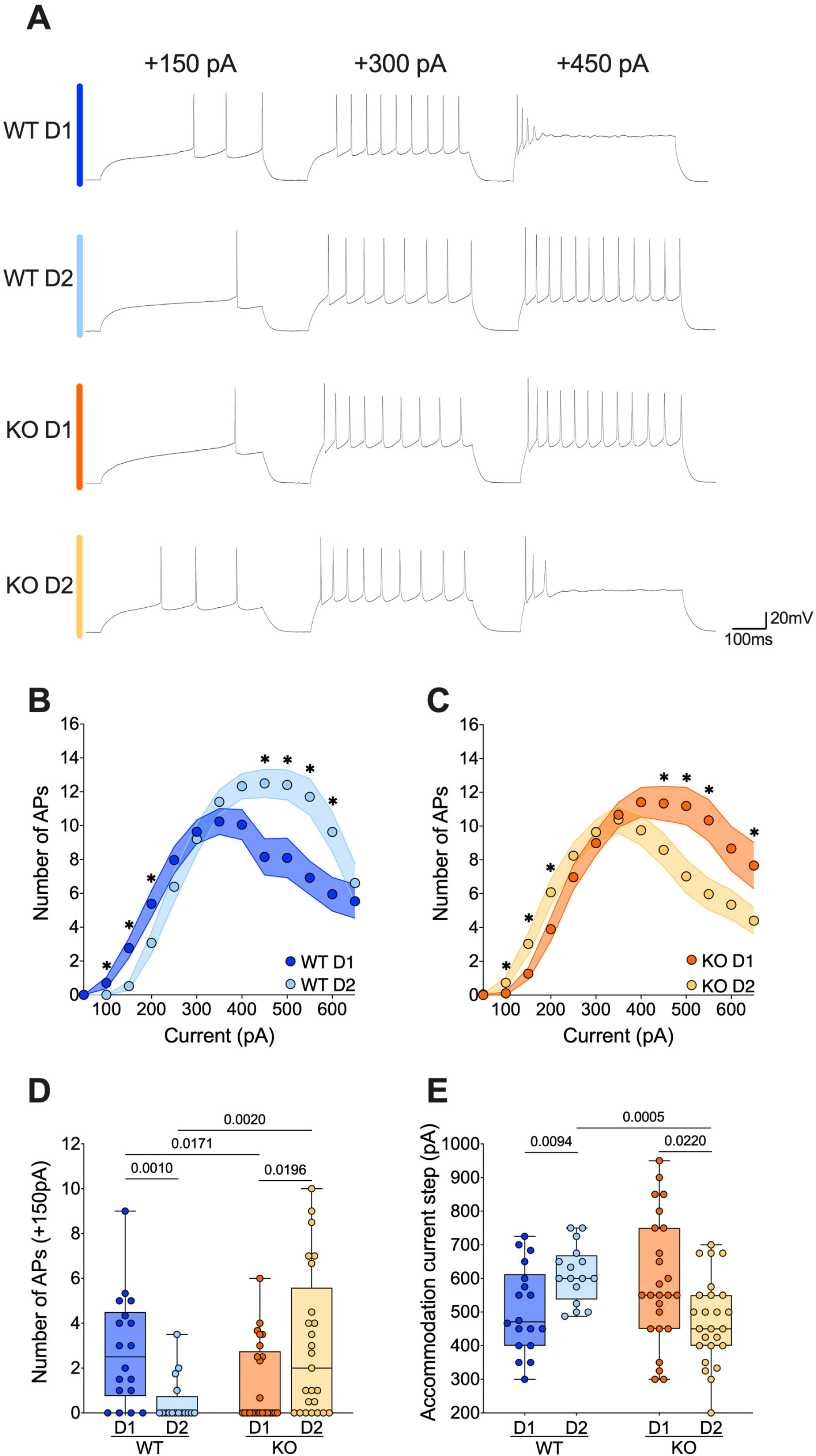
The active properties of accumbal D1- and D2-MSNs diverge in opposite ways in wild type and FXS mice. **(A)** Sample spike trains in response to depolarizing currents from D1- and D2-MSNs in each genotype. **(B, C)** Step by step analysis of the number of evoked action potentials in response to increasing depolarizing current shows that the hierarchy of excitability between D1- and D2-MSNs reverses in *Fmr1* KO mice. Single dot represents group mean value for each current step. Data are shown as mean ± SEM in XY plot. Multiple Mann-Whitney *U* test. * p-values <0.05. **(D)** At physiological depolarizing current steps, D1-MSNs are more excitable than D2-MSNs in WT mice, whereas in *Fmr1* KO mice, D1-MSNs are less excitable than D2-MSNs. **(E)** In WT mice the D1-MSNs accommodate their firing at lower current steps compared with D2-MSNs, while it is the exact opposite in *Fmr1* KO mice. **(D, E)** Single dot represents an individual mouse. Data are shown as min. to max. box plot with median and 25-75 percentile. Mann-Whitney *U* tests. P-values <0.05 are displayed in graphs. **(B-E)** WT D1-MSNs N=18 in dark blue, WT D2-MSNs N=16 in light blue, KO D1- MSNs N=25 in dark orange, KO D2-MSNs N=25 in light orange

### Divergent action potential properties of D1- and D2-MSNs in WT and FXS mice

Given the cell-specific impact of the absence of FMRP on the intrinsic excitability of MSNs, action potential kinetics were examined in our different cell types and genotypes. Analysis of AP duration **(Fig. 4A)** revealed that APs in D1-MSN were longer than that of D2-MSNs in WT, whereas an opposite dichotomy was found in *Fmr1* KO mice **(Fig. 4B)**. Specifically, ablation of *Fmr1* widened and shortened APs in D2- MSNs and D1-MSNs, respectively **(Fig. 4B)**. The loss of the normal dichotomy in AP duration was paralleled by opposite changes in AP amplitude of D1-MSNs and D2-MSNs in the absence of FMRP: compared to WT, APs got larger in D1- and smaller in D2-MSNs **(Fig. 4C)**. We reasoned that the observed differences in AP duration could result from phase specific changes in the depolarization and/or repolarization period **(Fig. 4D)**. In WTs, the AP depolarization phase was longer in D1- than in D2-MSNs; this physiological divergence disappeared in *Fmr1* KO mice **(Fig. 4E)**. AP repolarization times were similar across genotypes in D1-MSNs but not in D2-MSNs that repolarized slower in *Fmr1* KO mice **(Fig. 4F)**. Finally, comparing the AP undershoot phase, the so-called afterhyperpolarization (AHP) **(Fig. 4G)**, showed that while AHP amplitudes were similar in all groups **(Fig. 4H)**, AHPs were longer in D2-MSNs from *Fmr1* KO mice **(Fig. 4I)**.

**Figure 4.**
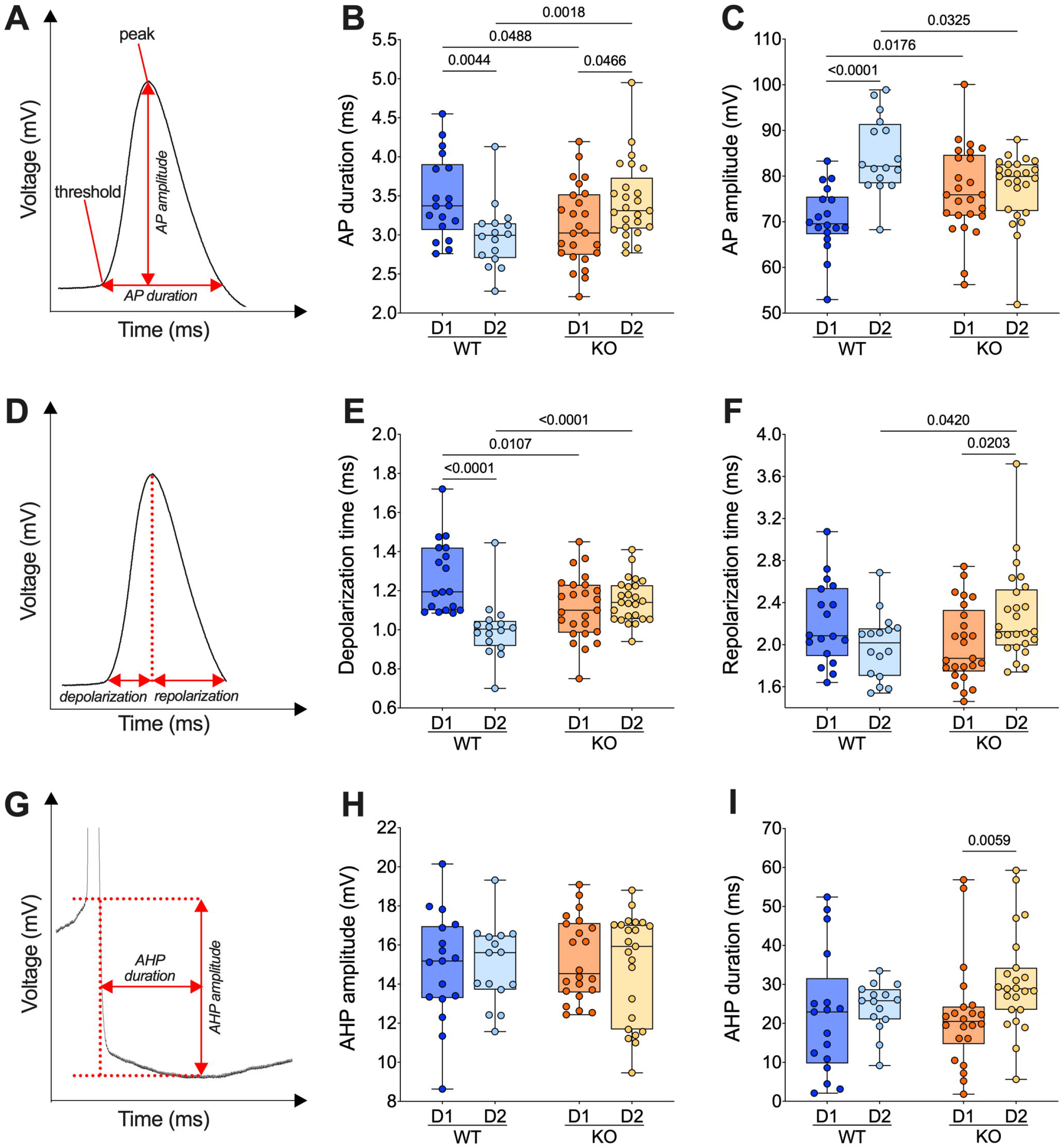
The action potential properties of D1- and D2-MSNs from WT mice present a dichotomy that is absent or reverted in FXS mice. **(A)** Example of individual AP evoked by depolarizing current injection, showing its threshold, peak, amplitude, and duration. **(B)** In WT mice, APs are shorter in D2- than D1-MSNs whereas in *Fmr1* KO mice, APs are longer in D2- than D1-MSNs. **(C)** In WT mice, APs are larger (i.e., the voltage difference between threshold and peak) in D2- than D1-MSNs but similar in *Fmr1* KO mice. **(D)** Example of individual AP evoked by depolarizing current injection, indicating depolarization and repolarization metrics. **(E)** AP depolarization time is shorter in D2- than in D1-MSNs in WT and similar in *Fmr1* KO mice. **(F)** The AP repolarization time of D2-MSNs is shorter in WT than *Fmr1* KO mice. **(G)** Example of individual AP evoked by depolarizing current injection, indicating AHP amplitude and AHP duration metrics. **(H-I)** AHP amplitude is comparable in all groups and its duration selectively increased in D2-MSNs from *Fmr1* KO mice. **(B, C, E, F, H, I)** Single dot represents an individual mouse. Data are shown as min. to max. box plot with median and 25-75 percentile. Mann-Whitney *U* tests. P-values <0.05 are displayed in graphs. **(B, C, E, F)** WT D1-MSNs N=18 in dark blue, WT D2-MSNs N=16 in light blue, KO D1-MSNs N=25 in dark orange, KO D2-MSNs N=25 in light orange. **(H-I)** WT D1-MSNs N=17 in dark blue, WT D2- MSNs N=15 in light blue, KO D1-MSNs N=22 in dark orange, KO D2-MSNs N=23 in light orange.

### The normal dichotomy that characterizes D1- and D2-MSNs is thrown out of balance in absence of FMRP

To summarize our comparative electrophysiological profiling, an index based on passive properties, excitability, and APs kinetics of these neurons was computed. The index reflected the strong dichotomy between D1- and D2-MSNs present in WTs. In *Fmr1* KO mice the dichotomy changed polarity, likely because of opposite adaptations in D1/D2-MSNs in the absence of FMRP **(Fig. 5)**.

**Figure 5.**
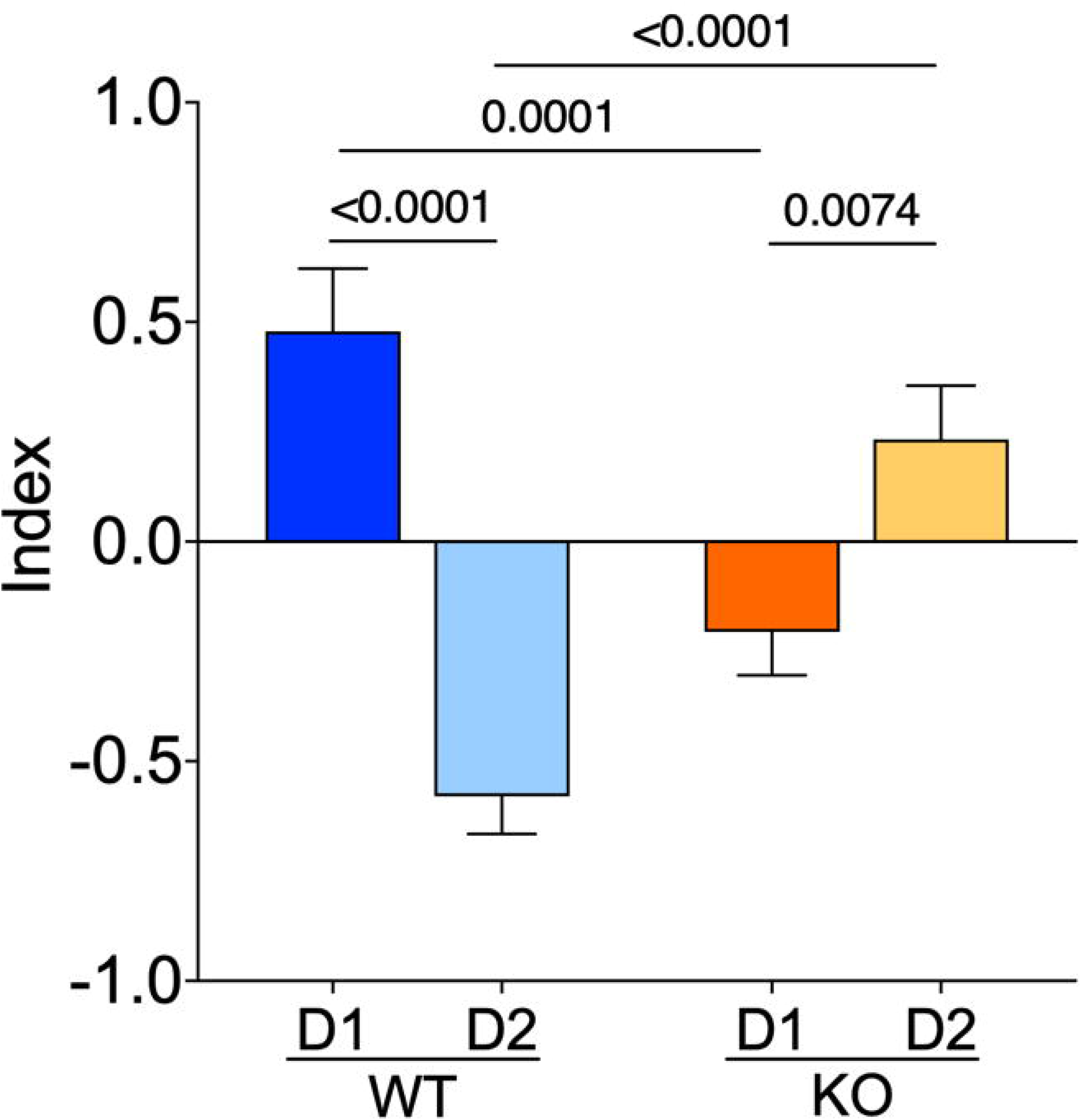
The normal dichotomy that characterizes D1- and D2-MSNs is thrown out of balance in absence of FMRP. D1- and D2-MSNs exhibit a dichotomy of their electrophysiological profile which is reversed in KO mice. FMRP’s absence impacts MSNs subtypes in opposite manners. The index combines input resistance, APs number at +150pA and APs duration parameters. Data were standardized per variable and plotted per group. Data are shown as mean with SEM column bar graph. Mann-Whitney *U* tests. P-values <0.05 are displayed in graphs. WT D1-MSNs N=18, WT D2-MSNs N=16, KO D1-MSNs N=25, KO D2-MSNs N=25 for each plotted variable.

### Multivariate analysis of the compound effects of *Fmr1* ablation in identified MSNs

Correlations among electrophysiological parameters have important functional implications in neurons. A multivariate analysis was used to evaluate how the lack of FMRP alters the relationships between pairs of electrophysiological features **(Fig. 6A, B, D, E)**. In WTs, MSN subtypes notably differed in sag-rheobase and accommodation-threshold correlations **(Fig. 6C)**. In contrast, *Fmr1* KO differed in “AP amplitude-latency” and “rheobase -input resistance” **(Fig. 6F)**. Across genotypes, D1-MSNs differed in AP duration-rheobase and AP amplitude-input resistance **(Fig. 6G)** while, D2-MSNs in AP amplitude-latency, AP amplitude-rheobase, AP number-rheobase, and accommodation-threshold **(Fig. 6H)**. These data further illustrate how FMRP deficiency strongly affects MSN subtypes’ electrophysiological profiles

**Figure 6.**
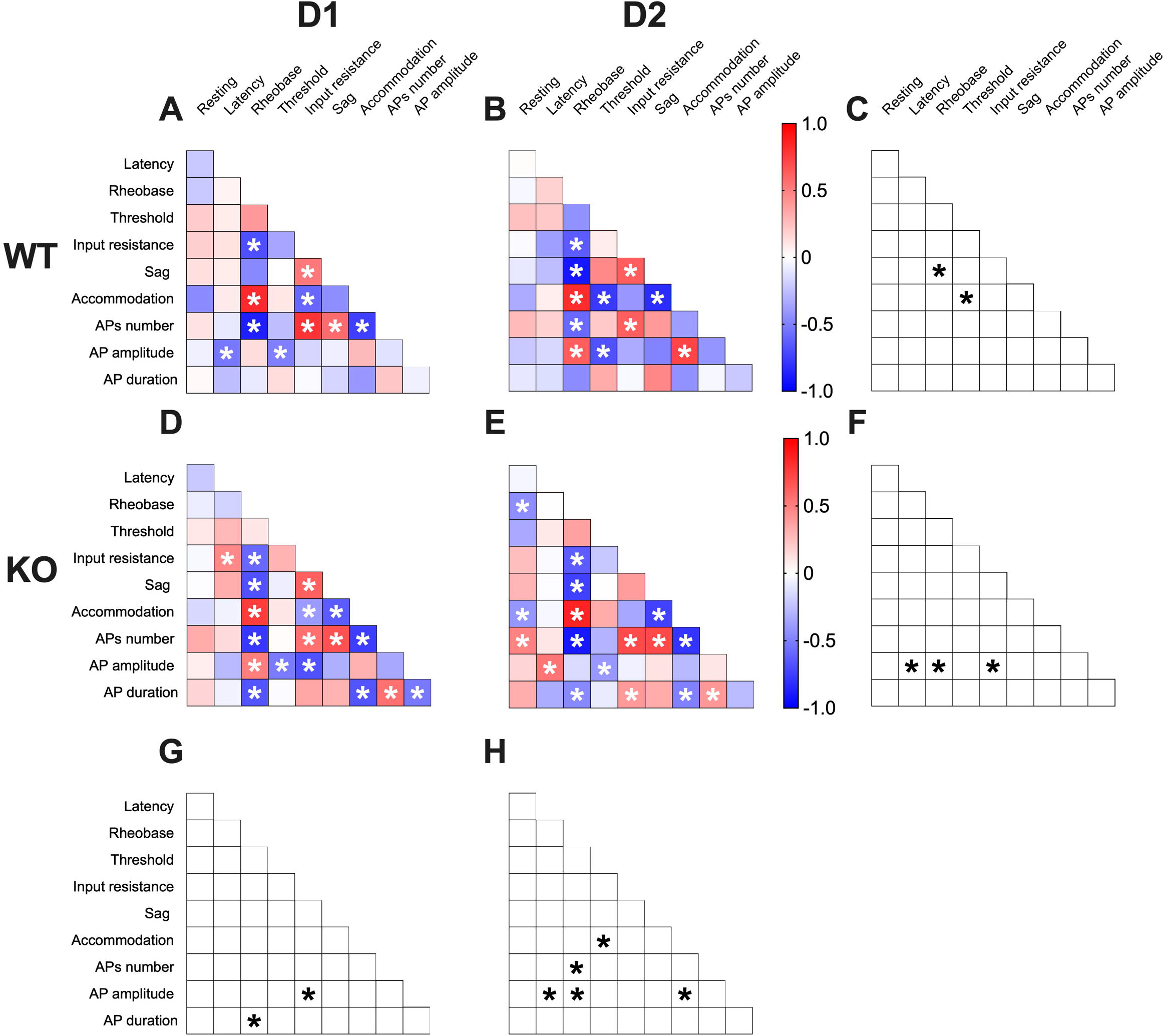
Multivariate analysis of the compound effects of *Fmr1* ablation in identified MSNs’ subtypes. **(A-B)** Heat maps of correlations’ profiles of electrophysiological parameters of D1- and D2- MSNs in WT or **(D-E)** *Fmr1* KO mice. **(A, B, D, E)** Non-parametric Spearman correlation matrix (r values). Statistically significant correlations (p-values <0.05) are displayed in graphs with white asterisk. **(C, F, G, H)** Correlations that significantly differ (p-value <0.05, Fisher1925) are indicated with a black asterisk by **(C, F)** genotype and **(G, H)** cell-type. **(A-H)** WT D1-MSNs N=18, WT D2-MSNs N=16, KO D1-MSNs N=25, KO D2-MSNs N=25.

## Discussion

FXS is the most common monogenic cause of autism and inherited intellectual disability [34]. Cognitive, emotional, and social deficits observed in FXS patients are indicative of dysfunction in the ventral striatum [19]. Although the striatum is an essential component of the autistic phenotype [35], studies have so far focused on the dorsal striatum, leaving the role of its ventral part poorly understood.

The present study provides the first cell-type specific electrophysiological profile of identified accumbal neurons in adult FXS mice. The data show that the functional dichotomy that characterizes WT MSN activity (i.e., D1-MSNs are more excitable than D2-MSNs), disappears in FXS. Thus, FMRP deficiency decreases the excitability of D1-MSNs and increases that of D2-MSNs. Mechanistically, this reversed phenotype is caused by both a more negative threshold and lower input resistance in D1-MSNs and a more depolarized resting potential, lower rheobase, and greater input membrane resistance in D2-MSNs.

Although the canonical separation of D1-direct and D2-indirect pathways in the accumbens is rightly disputed [36], the fact remains that D1-MSN and D2-MSN are major contributors to direct and indirect pathways, respectively [37]. Thus, the alteration in excitability between D1 and D2 reported here will alter accumbal outputs: the increased and decreased excitability of D2- and D1-MSNs, respectively, will likely unbalance mesolimbic pathways in favor of the indirect route. In keeping with this hypothesis, clinical and preclinical data both indicated a greater avoidance/approach behavior ratio in ASD, suggestive of an elevated tone of the indirect mesolimbic pathway [17].

In FXS, channel dysfunctions underlie pathological changes in neuronal excitability. FMRP regulates the gene expression or the function of various channels, thereby modulating AP properties [38]. Investigated AP kinetics in FXS mice showed that the absence of FMRP increased and decreased the amplitude of APs in D1- and D2-MSNs, respectively.

Additionally, AP duration was shorter in D1- and longer in D2-MSNs of FXS mice than in WT littermates. These changes in AP kinetics result from mirroring alterations in depolarization and repolarization phases, giving further credit to the idea that *Fmr1* ablation is linked to cell-specific modifications in the NAc. In D1-MSNs of FXS mice, repolarization was normal, but depolarization was shorter, in agreement with the known effects of FMRP ablation on voltage-gated Na^+^ channels [38]. D2-MSNs displayed longer depolarization and repolarization than WT. These changes are reminiscent to those reported in cortical and hippocampal neurons where the lack of FMRP influences big potassium (BK) channel conductance causing AP broadening and a higher firing probability [39]. Thus, a difference in BK activity could explain the altered repolarization observed in D2-MSNs.

Multivariate analysis uncovered covariation of selected intrinsic properties in a cell-specific manner in the absence of FMRP and suggested a degree of functional dependence in the associated neuronal properties. While some parameters covariate, keeping their correlation in all groups (e.g., rheobase-AP number), other parameters correlate differently across cell-types and genotypes (e.g., AP amplitude with latency, rheobase, input resistance and accommodation).

Dopaminergic control of accumbal D1 and D2 MSNs is central to reward behaviors [18]. DA D1R-mediated enhancement of L-type Ca^++^ channels augments D1-MSN excitability while binding of D2R has the opposite action in D2-MSNs [40–42]. FMRP is a key messenger for dopamine modulation in the striatum [43] and consequently dopamine signaling is altered in FXS mice [44]. It is tempting to hypothesize that the cell-type specific perturbation of excitability in D1- and D2-MSNs reported here results, at least partly, from dopaminergic dysfunctions in the NAc of FXS mice.

In conclusion, these results show that the absence of FMRP induces a cell-autonomous phenotype of MSN in the NAc and further emphasize the need to study the role of NAc in this disease.

## Abbreviations

ACSF: Artificial cerebrospinal fluid
AHP: Afterhyperpolarization
AP: Action potential
BK: Big potassium
D1R: dopamine receptor type 1
D2R: dopamine receptor type 2
*Drd1*: Dopamine receptor D1 gene
*Drd2*: Dopamine receptor D2 gene
EGFP: Enhanced green fluorescent protein
*Fmr1*: Fragile X mental retardation 1 gene
FMRP: Fragile X mental retardation protein
FXS: Fragile X syndrome
ISH: in situ hybridization
KO: Knockout
MSN: Medium spiny neuron
NAc: Nucleus accumbens
smFISH: Single molecule fluorescent in situ hybridization
WT: Wild type

## Declarations

### Ethics approval and consent to participate

Animals were treated in compliance with the European Communities Council Directive (86/609/EEC) and the United States National Institutes of Health Guide for the care and use of laboratory animals. The French Ethical committee authorized this project (APAFIS#3279-2015121715284829 v5).

### Authors’ contributions

GG: Conceptualization, Data curation, Formal analysis, Validation, Writing—review and editing.

OL: Data curation, Formal analysis, Validation, Methodology. LMM: Data curation, Formal analysis, Validation.

EV: Data curation, Formal analysis, Validation, Writing—review and editing. PC: Conceptualization, Supervision, Methodology, Writing—review and editing

OJJM: Conceptualization, Supervision, Funding acquisition, Methodology, Project administration, Writing— original draft, review and editing.

## Acknowledgements

The authors are grateful to the Chavis-Manzoni team members for helpful discussions and to Dr. A.F. Scheyer for critical reading and help with writing the manuscript.

## Supplementary Figures Legends

**Supplementary Figure 1. The absence of FMRP induces cell-specific alterations in the membrane voltage response to hyperpolarizing current injections.**

**(A)** Examples of membrane voltage response to hyperpolarizing current injections in our experimental groups. **(B)** In response to hyperpolarizing current injections D1-MSNs membrane voltage response is greater in WT than *Fmr1* KO mice. **(C)** On contrary, D2- MSNs show a lower response in WT than *Fmr1* KO mice. **(D)** Accordingly, in absence of FMRP, D1-MSNs have a higher input resistance, **(E)** while D2-MSNs have a lower one. **(B-E)** Single dot represents group mean value at that current step. Data are shown as mean ± SEM in XY plot. Multiple Mann-Whitney *U* test. ⍰ p-values <0.05. **(F)** Example of membrane voltage response to a -400pA current injection, indicating the sag voltage. **(G)** In response to a -400pA current injection D2-MSNs exhibit a smaller sag voltage in WT compare with FXS mice. Single dot represents an individual mouse. Data are shown as min. to max. box plot with median and 25-75 percentile. Mann-Whitney *U* tests. P-values <0.05 are displayed in graphs. **(B-E, G)** WT D1-MSNs N=18 in dark blue, WT D2-MSNs N=16 in light blue, KO D1- MSNs N=25 in dark orange, KO D2-MSNs N=25 in light orange.

**Supplementary Figure 2. In absence of FMRP D1- and D2-MSNs excitability change in opposite.**

**(A)** Example of firing pattern triggered by the injection of +150pA, +300pA and +450pA depolarizing currents for each group. **(B, C)** Analysis by cell-type of the number of evoked action potentials in response to increasing depolarizing current shows that FMRP absence has an opposite effect on D1- and D2-MSNs firing profile. Single dot represents group mean value at that current step. Data are shown as mean ± SEM in XY plot. Multiple Mann-Whitney *U* test. ⍰ p-values <0.05. **(D)** No statistically significant difference was found in the maximum firing rate achieved by MSNs during increasing steps of depolarizing currents. **(E)** Analysis of the amount of current required to induce the higher firing rate showed that in WT mice D2-MSNs reached their maximum firing at higher current steps than D1-MSNs, whereas it was the opposite between MSNs of FXS mice. **(D, E)** Single dot represents an individual mouse. Data are shown as min. to max. box plot with median and 25-75 percentile. Mann-Whitney *U* tests. P-values <0.05 are displayed in graphs. **(B-E)** WT D1- MSNs N=18 in dark blue, WT D2-MSNs N=16 in light blue, KO D1-MSNs N=25 in dark orange, KO D2-MSNs N=25 in light orange.

