## Supplementary figures and images for "Dichotomous Intrinsic Properties of Adult Accumbens Medium Spiny Neurons Vanish in the Fragile X Mouse Model of Autism"

### Supplemental Figure 1

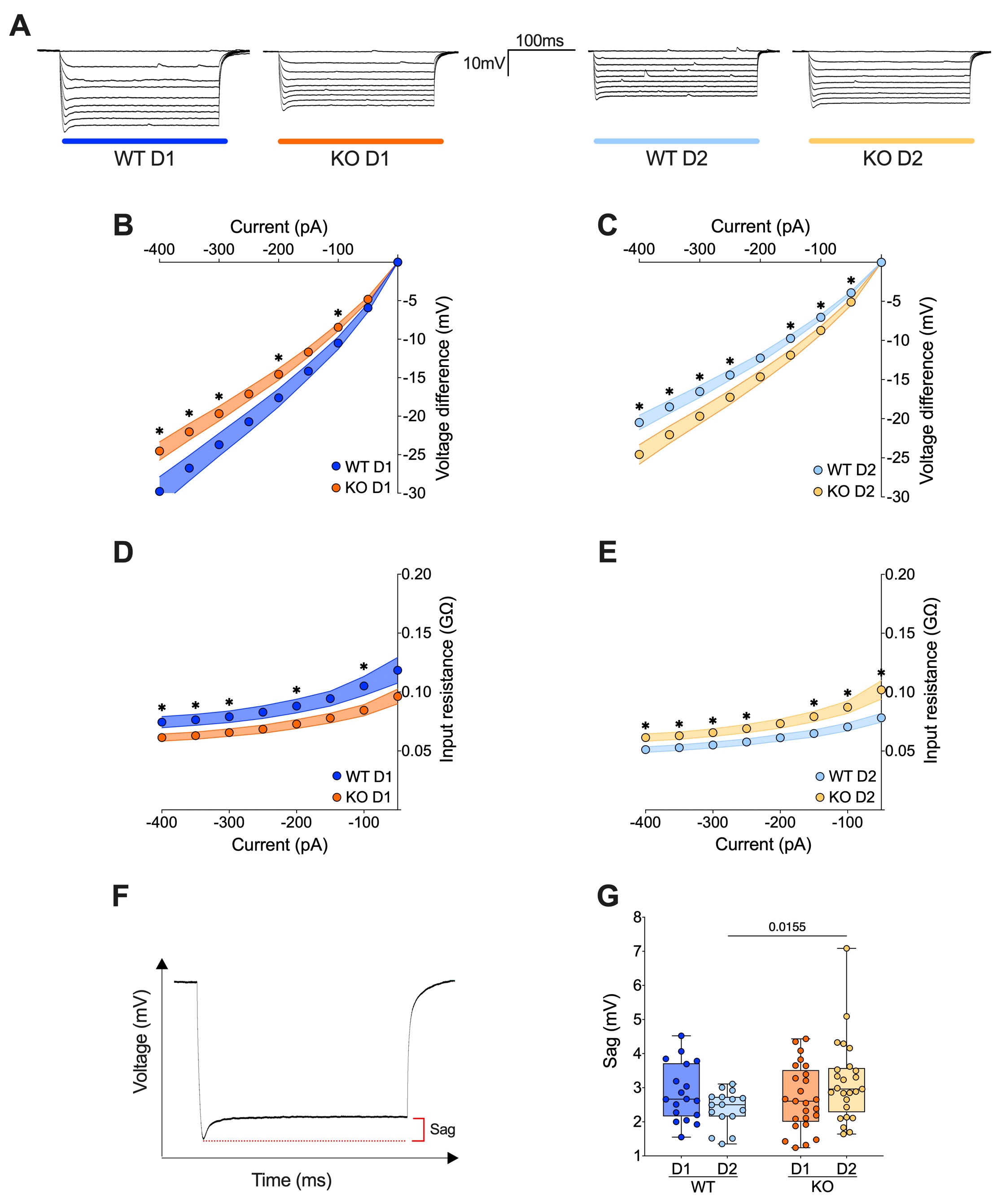

### Supplemental Figure 2

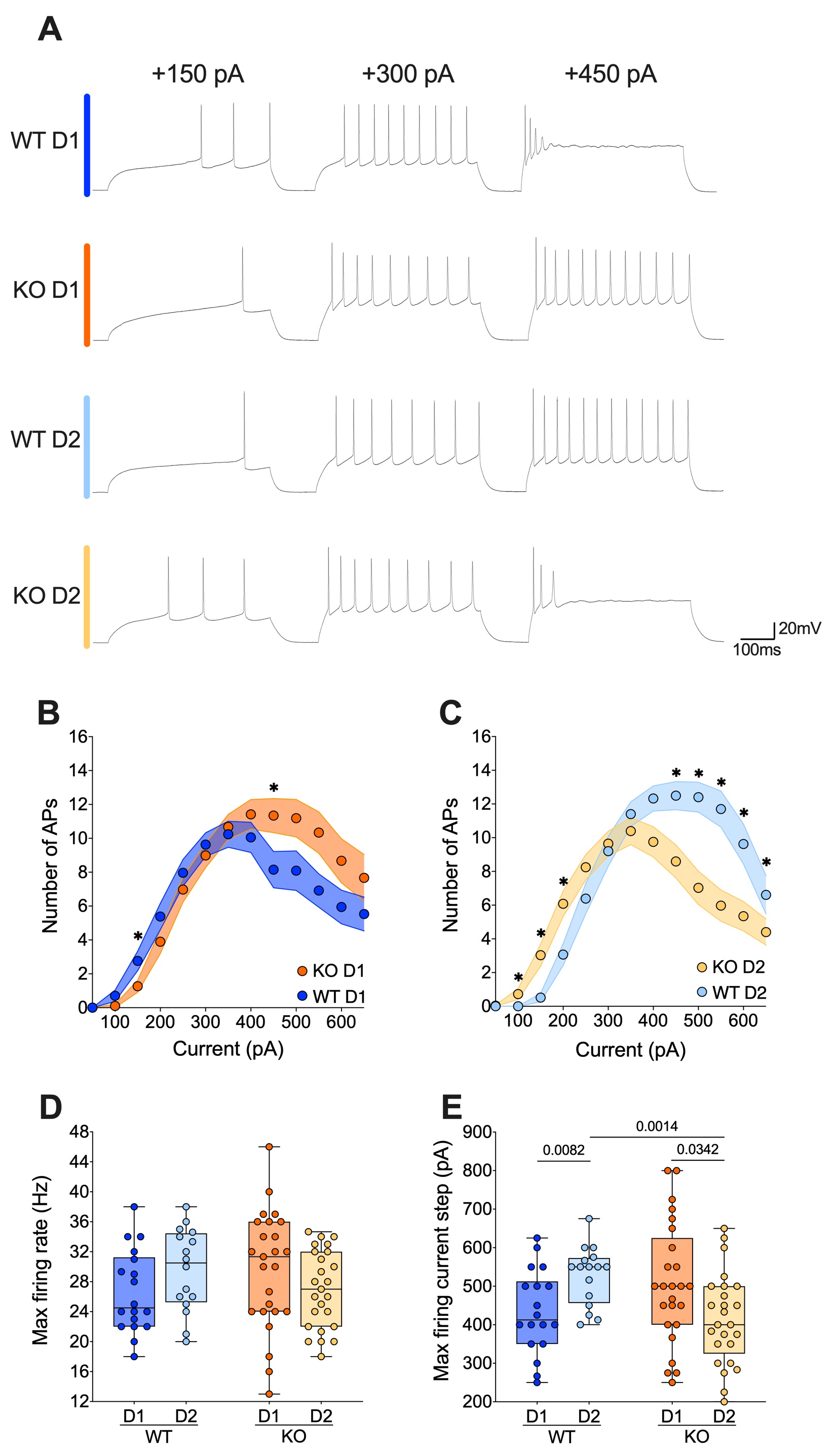
